# Comparative immunogenicity and protective efficacy of formaldehyde-inactivated whole-cell and sonicated whole-cell extract vaccines against multidrug-resistant *Pseudomonas aeruginosa* in mice

**DOI:** 10.64898/2026.09.09.750315

**Authors:** Rubaiya Binte Kabir, Jubayer Ahmed, Mahmuda Yasmin, SM Shamsuzzaman, Chowdhury Rafiqul Ahsan

**Affiliations:** Department of Microbiology, Dhaka Medical College, Dhaka, Bangladesh; Department of Microbiology, University of Dhaka, Dhaka, Bangladesh

**Keywords:** *Pseudomonas aeruginosa*, multidrug resistance, vaccine, whole-cell vaccine, sonicated whole cell extract, immunogenicity, serum bactericidal assay, mice

## Abstract

**Background:** *Pseudomonas aeruginosa* is a major opportunistic Gram-negative pathogen responsible for healthcare-associated infections, while increasing antimicrobial resistance has narrowed therapeutic options. Vaccine development therefore represents an important complementary strategy for prevention. This study compared the immunogenicity and protective efficacy of a formaldehyde-inactivated whole-cell vaccine with a sonicated whole cell extract preparation against multidrug-resistant (MDR) *P. aeruginosa* in mouse model.

**Methods:** Clinical *P. aeruginosa* isolates were obtained from specimens collected at Dhaka Medical College Hospital between July 2022 and February 2026. Formaldehyde-inactivated whole-cell and sonicated whole cell extract preparations were produced from *P. aeruginosa*. A total of 144 female Swiss albino mice aged 6–8 weeks were used: 120 mice were allocated to two independent immunization experiments and 24 mice were used for LD50 determination. Mice received three intramuscular immunizations on days 0, 14, and 28. Antigen-specific serum IgG was assessed by enzyme-linked immunosorbent assay (ELISA), functional antibody activity was evaluated by serum bactericidal assay (SBA), and protective efficacy was assessed following lethal intraperitoneal challenge.

**Results:** Ten isolates from MDR *P. aeruginosa* were selected by simple random sampling from the clinical isolates for vaccine preparation. The estimated protein concentration of the sonicated preparation was 1.872 mg/mL. Both vaccine groups developed significantly higher anti-*P. aeruginosa* IgG responses than the control group after immunization (p<0.001). Following lethal challenge, survival was 100% in mice receiving the formaldehyde-inactivated vaccine and 91.7% among mice receiving the sonicated preparation; no control mice survived. Between-vaccine differences in survival were not statistically significant. SBA demonstrated complement-dependent bactericidal activity in sera from both vaccinated groups, with activity detectable to a dilution of 1:32 for the whole-cell vaccine and 1:64 for the sonicated preparation.

**Conclusions:** Both vaccine preparations elicited strong humoral responses and substantial protection against lethal MDR *P. aeruginosa* challenge in mice. The findings support further investigation of multicomponent *P. aeruginosa* vaccine strategies, although additional studies using multiple challenge strains and broader immunological endpoints are required before conclusions regarding translational efficacy can be drawn.

## Introduction

*Pseudomonas aeruginosa* is a ubiquitous opportunistic Gram-negative pathogen and an important cause of healthcare-associated infection, particularly among hospitalized and immunocompromised patients. It is associated with pneumonia, bacteremia, urinary tract infection, wound and surgical-site infection, and chronic pulmonary disease. Its capacity to persist in diverse environments, form biofilms, and acquire resistance to multiple antimicrobial classes makes treatment increasingly difficult (1,2). Carbapenem-resistant *P. aeruginosa* has been recognized as a critical-priority pathogen requiring new preventive and therapeutic approaches (3).

The continuing emergence of antimicrobial resistance is associated with prolonged hospitalization, increased healthcare expenditure, and poorer clinical outcomes (4). Although development of new antimicrobial agents remains necessary, vaccination could complement antimicrobial therapy by preventing infection, reducing antibiotic exposure, and potentially limiting selection and transmission of resistant organisms (5,6). Despite extensive investigation of candidate antigens, no licensed *P. aeruginosa* vaccine is currently available.

Whole-cell inactivated vaccines are attractive because they present a broad repertoire of bacterial antigens and pathogen-associated molecular patterns. Chemical inactivation can eliminate bacterial infectivity while retaining antigenic structures. In contrast, protein-based preparations can focus immune responses on antigenic components while potentially reducing some of the reactogenicity associated with intact bacterial cells. Outer membrane and other membrane-associated proteins are of particular interest because several are conserved and immunogenic across clinical isolates.

Sonication disrupts bacterial cells and generates a complex protein-containing preparation that may include membrane-associated and soluble antigens. Such multicomponent preparations could broaden immune recognition in the face of the substantial antigenic diversity of *P. aeruginosa*. The protective value of a vaccine candidate, however, should not be inferred solely from antibody concentration; functional antibody activity and survival after challenge provide complementary evidence. ELISA measures antigen-specific antibody responses, whereas SBA assesses complement-dependent bactericidal activity.

Previous studies have investigated LPS, exotoxin A, flagellar proteins, alginate, outer membrane proteins, and type III secretion system components, but none of them has yet achieved sufficient breadth and efficacy for routine clinical use against *P. aeruginosa* (Killough et al., 2022). Evidence evaluating vaccine candidates against MDR *P. aeruginosa* in Bangladesh remains limited. Therefore, this study compared the immunogenicity and protective efficacy of formaldehyde-inactivated whole-cell and sonicated whole cell extract preparations in mouse model.

## Materials and Methods

### Study design and setting

This experimental study was conducted at the Departments of Microbiology, Dhaka Medical College and the University of Dhaka, Bangladesh, from July 2022 to February 2026. *P. aeruginosa* was collected from clinical specimens obtained from patients attending Dhaka Medical College Hospital using purposive sampling. A total of 10 MDR *P. aeruginosa* were selected by simple random sampling from the stock culture further for vaccine preparation.

### Antimicrobial susceptibility testing

Antimicrobial susceptibility testing was performed using the Kirby–Bauer disk diffusion method on Mueller–Hinton agar in accordance with the Clinical and Laboratory Standards Institute (CLSI) guidelines (7). *P. aeruginosa* ATCC 27853 was used as the quality control strain.

The antimicrobial agents tested were piperacillin (100 µg), piperacillin/tazobactam (100/10 µg), ceftazidime (30 µg), aztreonam (30 µg), imipenem (10 µg), meropenem (10 µg), amikacin (30 µg), gentamicin (10 µg), netilmicin (30 µg), ciprofloxacin (5 µg), levofloxacin (5 µg), and doxycycline (30 µg) (Oxoid Ltd., UK). Bacterial suspensions equivalent to a 0.5 McFarland standard were inoculated onto Mueller–Hinton agar plates, antibiotic discs were applied, and the plates were incubated aerobically at 37°C for 16–18 hours. The inhibition zone diameters were interpreted according to CLSI breakpoints (7). The minimum inhibitory concentration (MIC) of colistin was determined by the agar dilution method following CLSI recommendations (7). Acquired non-susceptibility to at least one agent in three or more antimicrobial categories (such as, amikacin or gentamycin, piperacillin-tazobactam imipenem or meropenem, ceftazidime or cefepime, ciprofloxacin or levofloxacin, colistin (when no other option)) was considered as multidrug resistance (8).

### Experimental animals and allocation

A total of 144 female Swiss albino mice aged 6–8 weeks were collected from ICDDR,B. Sixty mice were allocated to each of two independent immunization experiments, with three groups in each experiment containing 20 mice in each group: Group 1, formaldehyde-inactivated whole-cell vaccine; Group 2, sonicated protein preparation; and Group 3, PBS control. Twelve mice per group were used for lethal challenge and 8 mice per group were retained for post-challenge serum assessment. Twenty-four additional mice were used for LD50 determination. They were kept in the animal house of the Department of Microbiology, Dhaka Medical College. Well ventilated cages, optimum temperature of 25°C, proper light and dark cycle, standard laboratory pellet feeds and clean water ad libitum throughout the experimental period had been maintained. Mice were given acclimatization period of two weeks before starting the experiments. They were closely monitored for any clinical signs of distress or deterioration predefined as humane endpoints.

### Preparation of formaldehyde-inactivated whole-cell vaccine

*P. aeruginosa* was cultured in tryptic soy broth and incubated at 37°C following growth on Mueller– Hinton agar. The culture was centrifuged (2,000 rpm, 20 min, 4°C) and the pellet was washed twice with PBS. Formalin (3%) was added, followed by incubation at 37°C for 2 h. The preparation was then washed twice with PBS, adjusted to 1.5×10^8 CFU/mL, and diluted to 2×10^7 CFU/mL. Complete inactivation was confirmed by absence of bacterial growth on Mueller–Hinton agar. Supernatant containing the whole cell extract was stored at −20°C until used (9).

### Preparation of sonicated whole cell extract

A loopful of *P. aeruginosa* was inoculated into 100 mL tryptic soy broth and incubated with shaking at 37°C. Forty millilitres of culture were centrifuged at 8,000 rpm for 10 min. The pellet was washed with PBS, resuspended in 3 mL distilled water, and sonicated at 20 kHz for two 10-s pulses while maintained on ice. The lysate was centrifuged at 8,000 rpm for 8 min at 4°C, and the supernatant was collected and stored at −20°C until used (10).

### Protein quantification

Protein concentration of the sonicated whole cell extract was measured. The sonicated preparation had an absorbance of 1.523 at 595 nm, corresponding to an estimated protein concentration of 1.872 mg/mL.

### Immunization

Mice received three intramuscular immunizations using a 31-gauge insulin syringe on day 0, 14, and 28. Group 1 received 20 µL of the formaldehyde-inactivated preparation, Group 2 received 20 µL of the sonicated preparation containing approximately 3μg of protein, and Group 3 received 20 µL PBS (9).

### Serum collection

Tail blood was collected 14 days after each immunization (days 14, 28, and 42) from 8 mice per group for ELISA. Blood was diluted in PBS (50μl in 200μl PBS to yield a dilution of 1:5), centrifuged at 3,000 rpm for 10 min, and the recovered serum was stored at −20°C until analysis. Proper anesthesia with intra-peritoneal injection of Ketamine (dose 100 mg/kg) was used before tail blood collection (9).

### LD50 determination and challenge

Twenty-four mice were divided into four groups of six and challenged with serial concentrations of *P. aeruginosa* corresponding to approximately 10^6–10^9 CFU/mL. Mortality was monitored, and the dose producing approximately 50% mortality was designated the LD50. In the reported experiment, the 10^8 CFU/mL preparation resulted in 4/6 deaths (66.7%) and was used as the LD50 dose. Two weeks after the final immunization, 12 mice from each group were challenged intraperitoneally with 100 µL of a bacterial suspension described as three times the LD50 (5). The challenge strain of *P. aeruginosa* was MDR which was resistant to piperacillin, piperacillin-tazobactam, ceftazidime, and imipenem. Animals were observed for clinical signs and mortality for 30 days after challenge.

### Serum collection by cardiac puncture

After 1 month of challenge, blood collection by cardiac puncture was done from rest of the remaining 8 mice (unchallenged) from both experimental and control groups for detection of serum IgG titre by ELISA and evaluation of functional capacity of the immunized serum by Serum Bactericidal Assay (SBA) (10).

### ELISA

After sonication of whole cell *P. aeruginosa,* it was diluted to 10 µg/mL in carbonate–bicarbonate buffer (pH 9.6) and considered as ideal quality. After coating microtiter plate, they were washed with PBS 0.05% Tween-20 and blocked with 5% skim milk in PBS Tween-20. Mice sera was used as primary antibody. Horseradish peroxidase-conjugated anti-mouse IgG (1:5,000) was used as the secondary antibody. Tetramethylbenzidine/urea peroxide substrate was used for color development, and the reaction was stopped with 1 M sulfuric acid. Absorbance was measured at 450 nm. The cutoff was defined as the mean OD of the negative-control sera plus two standard deviations.

### Serum bactericidal assay (SBA)

SBA was performed with modifications of the method described by Son and Taylor in 2011 (11). An overnight *P. aeruginosa* culture was washed and adjusted to an OD600 of 0.1, corresponding to approximately 10^8 CFU/mL, followed by dilution to the working inoculum. Guinea pig complement was used as the exogenous complement source. Two-fold serial dilutions of immune and control sera were prepared in PBS. Serum, bacterial suspension, and complement were combined and incubated at 37°C for 1 h. Tryptic soy broth was then added, followed by incubation for 2 h. Aliquots were plated on MacConkey agar and colonies were counted after overnight incubation. Bactericidal activity was assessed by comparison with control sera. Counts reported as too numerous to count (TNTC) were recorded as such.

### Statistical analysis

Data were entered and analyzed using IBM SPSS Statistics version 27.0, Microsoft Excel, and R Studio version 2026.08.2+200. Continuous variables were summarized using means and standard deviations, and categorical variables using frequencies and percentages. Differences in ELISA optical-density values across immunization time points were evaluated using repeated measure ANOVA, followed by post-hoc comparisons with Bonferroni adjustment where applicable. Survival between groups was assessed using chi-square and Fisher’s exact tests as reported. A two-sided p value <0.05 was considered statistically significant.

### Ethics approval

This study was approved by the Ethical Review Committee and Animal experimentation Ethics committee of Dhaka Medical College (Memo no: ERC-DMC/ERC/2022/156). Informed written consent were obtained from the participants. Animal experimentation was complied with ARRIVE 2.0 guideline. A complete ARRIVE checklist has been provided as Supporting information (S1 ARRIVE 2.0 checklist.

## Results

### Antigen-specific IgG responses

All the serum sample from experimental groups showed IgG absorbance values above the cut off value of 0.19, after 14 days of 1^st^ dose, 0.22 after 14 days of 2^nd^ and 3^rd^ dose. Following the first immunization, the mean OD450 was 0.37±0.017 in both vaccine groups, compared with 0.16 in the control group (Fig 1a). After the second immunization, mean OD450 value increased to approximately 1.18±0.03 in both vaccine groups, compared with 0.16 in controls (Fig 1b). After 3^rd^ dose, mean OD450 value was 1.19±0.05 in both experimental groups, compared to 0.12 in control group (Fig 1c). The reported difference between vaccinated and control groups was statistically significant (p<0.05).

**Fig 1:**
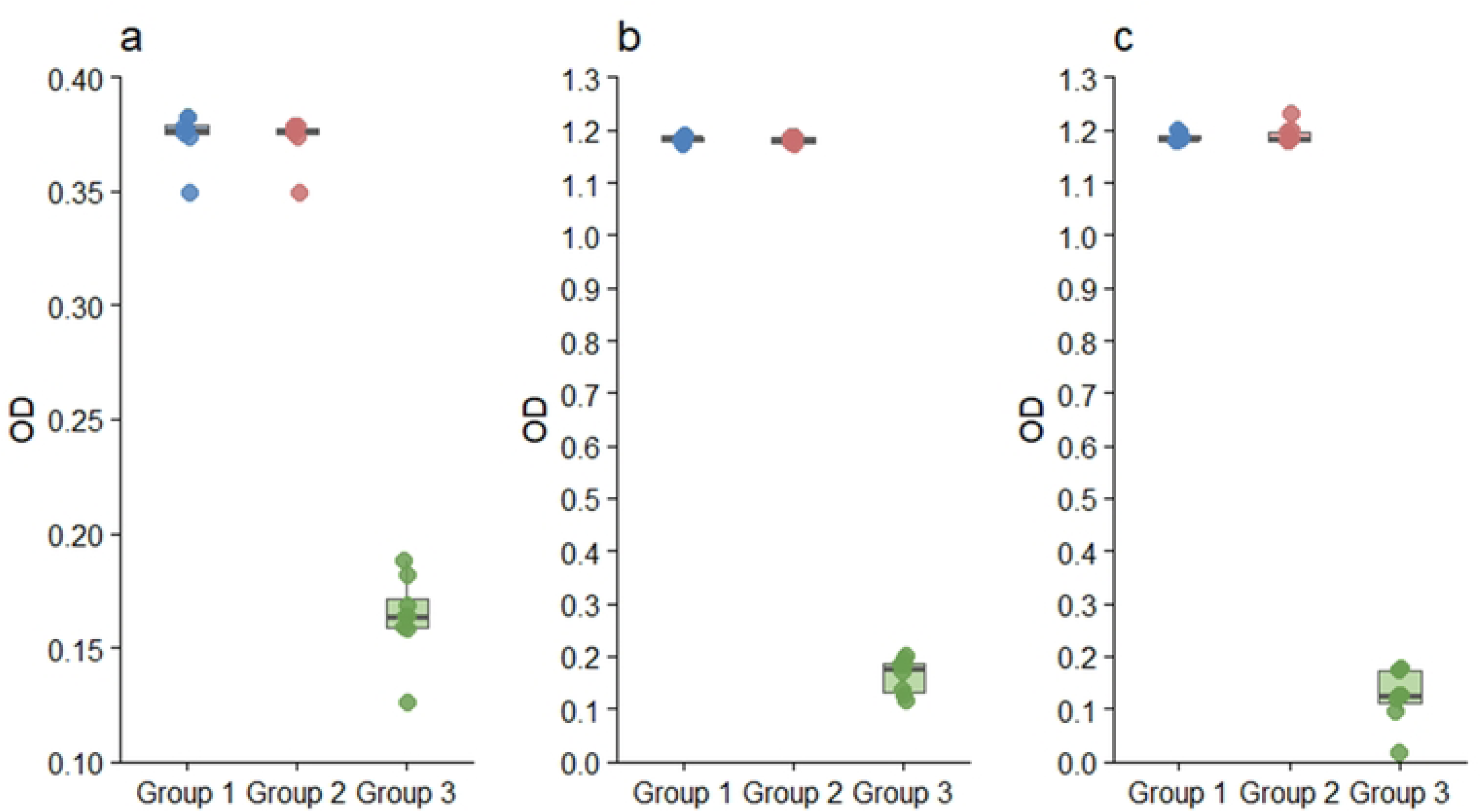
Serum antibody responses in mice following successive immunizations with formaldehyde-inactivated and sonicated *Pseudomonas aeruginosa* vaccines. (a) First immunization, (b) second immunization, and (c) third immunization. Each box represents the interquartile range with the median indicated by the horizontal line; individual points represent individual mice (n = 8).

In case of formaldehyde inactivated vaccine, a one-way repeated-measures ANOVA showed a significant effect of dose on antibody response, F(2, 14) = 48,034.91, p < 0.001. The antibody response increased markedly following the second dose and remained elevated following the third dose, with no significant difference between the second and third doses after Bonferroni correction (Table 1).

**Table 1:** Repeated-measures ANOVA and Bonferroni post-hoc analysis of antibody responses following successive doses of formaldehyde-inactivated vaccine.

| Analysis | Comparison | Test statistic | df | Adjusted p value | Interpretation |
| --- | --- | --- | --- | --- | --- |
| Repeated measure ANOVA | Dose | F = 48,034.91 | 2,14 | <0.001 | Significant |
| Bonferroni post- hoc | 1<sup>st</sup> vs 2<sup>nd</sup> dose | t = -271.0 | 7 | $7.45 \times 10^{-15}$ | Significant |
| Bonferroni post- hoc | 1<sup>st</sup> vs 3<sup>rd</sup> dose | t = -208.0 | 7 | $4.66 \times 10^{-14}$ | Significant |
| Bonferroni post- hoc | 2<sup>nd</sup> vs 3<sup>rd</sup> dose | t = -2.22 | 7 | 0.186 | Not Significant |

A similar pattern was observed for the sonicated preparation, F(2,14) = 16,964.62, p < 0.001 (Table 2).

**Table 2:** Repeated-measures ANOVA and Bonferroni post-hoc analysis of antibody responses following successive doses of sonicated antigens vaccine.

| Analysis | Comparison | Test statistic | df | Adjusted p value | Interpretation |
| --- | --- | --- | --- | --- | --- |
| Repeated measure ANOVA | Dose | F = 16,964.62 | 2,14 | <0.001 | Significant |
| Bonferroni post- hoc | 1<sup>st</sup> vs 2<sup>nd</sup> dose | t = -248.0 | 7 | $1.38 \times 10^{-14}$ | Significant |
| Bonferroni post- hoc | 1<sup>st</sup> vs 3<sup>rd</sup> dose | t = -129.0 | 7 | $1.31 \times 10^{-12}$ | Significant |
| Bonferroni post- hoc | 2<sup>nd</sup> vs 3<sup>rd</sup> dose | t = -2.04 | 7 | 0.242 | Not Significant |

### LD50 and survival following challenge

The reported LD50 experiment showed 66.7% mortality at 10^8 CFU/mL (Table 3); the selected lethal dose was considered three times the concentration of 10^8 CFU/mL.

**Table 3:** LD_50_ determining concentration of bacterial solution after 30 days of injecting mice:

| Bacterial concentration (CFU/mL) | Survived, n (%) | Died, n (%) | Total |
| --- | --- | --- | --- |
| $10^6$ | 6 (100) | 0 (0) | 6 |
| $10^7$ | 5 (83.3) | 1 (16.7) | 6 |
| $10^8$ | 2 (33.3) | 4 (66.7) | 6 |
| $10^9$ | 0 (0) | 6 (100) | 6 |

All mice receiving the formaldehyde-inactivated vaccine survived lethal challenge. Survival among mice receiving the sonicated preparation was 91.7%. No control animals survived (Fig 2). Overall comparisons between the three groups were statistically significant χ²≈32.05, p<0.0001). Pairwise Fisher’s exact testing showed significantly higher survival in both vaccinated groups than in controls, whereas the difference between the two vaccine groups was not statistically significant. Duplicate experiment presented similar result.

**Fig 2:**
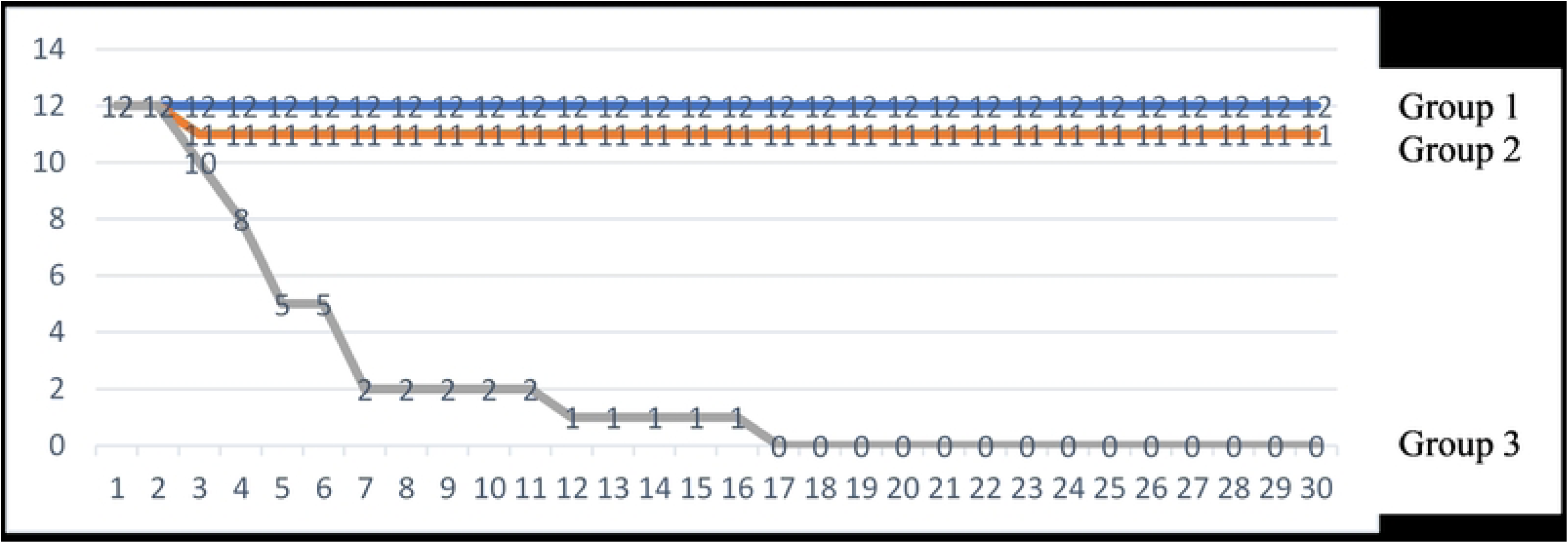
Graph showing survival of mice among 3 groups over 1 month after lethal challenge All mice of group 1 survived. Only one mouse died on the 3^rd^ day; the rest of the mice survived the lethal challenge in group 2. In the control group, two died on the 3^rd^ day, two more died on the 4^th^ day, three died on the 5^th^ day, three died on the 7^th^ day, one died on the 12^th^ day, last one died on the 17^th^ day.

### Post-challenge IgG response

Serum collected by cardiac puncture after the challenge phase showed mean OD450 values of approximately 1.17±0.03 in the formaldehyde-inactivated group and 1.16±0.03 in the sonicated group (both of which are above the cut off value of 0.21), compared with 0.15 in controls. The reported difference between vaccinated and control groups was statistically significant (p<0.05) (Fig 3).

**Fig 3:**
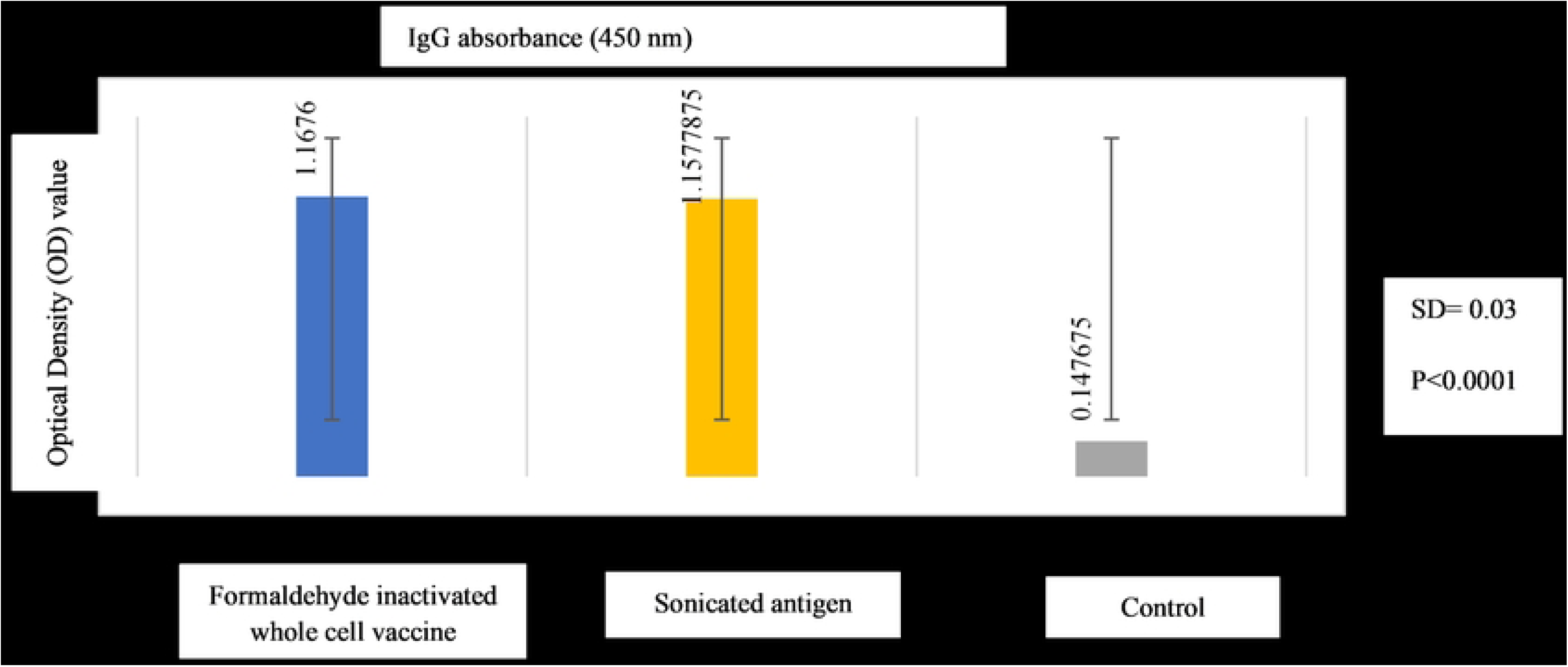
OD450 values of serum IgG absorbance after cardiac puncture by ELISA (SD= 0.03, p<0.05) (Bars represent mean OD and error bars represent standard deviation (SD))

### Serum bactericidal activity

Sera from both vaccinated groups demonstrated complement-dependent bactericidal activity at low serum dilutions. Activity was detectable through 1:32 in the whole-cell vaccine group and through 1:64 in the sonicated vaccine group (Fig 4). These findings descriptively suggest greater bactericidal activity of serum from the sonicated-vaccine group.

**Figure 4:**
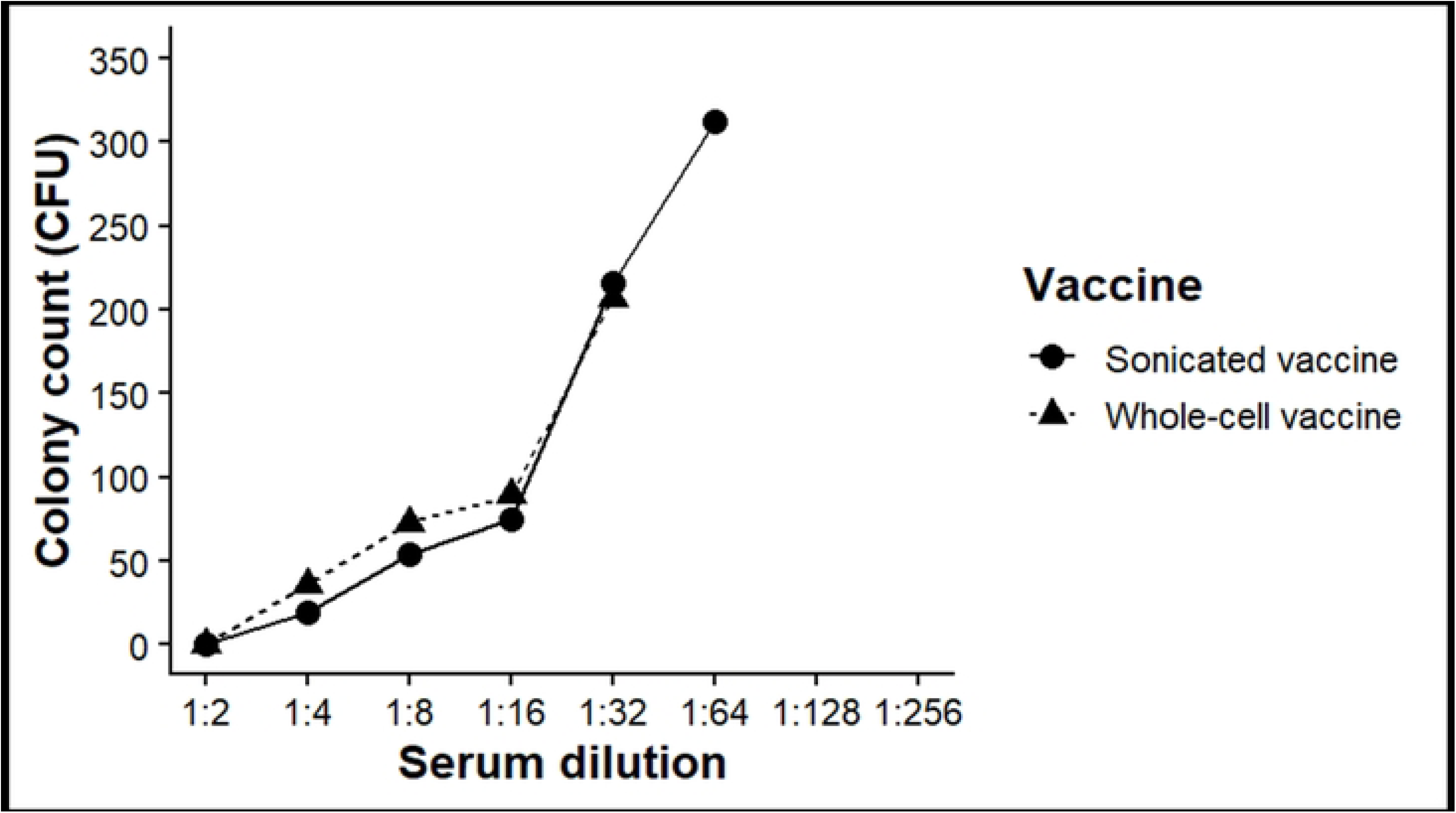
Serum bactericidal activity of whole-cell and sonicated *P. aeruginosa* vaccine preparations.

## Discussion

The present study evaluated two multicomponent *P. aeruginosa* vaccine preparations— formaldehyde-inactivated whole cells and a sonicated protein preparation—in a murine model. Both formulations generated strong antigen-specific IgG responses, functional serum bactericidal activity, and substantial protection against lethal intraperitoneal challenge with the study strain. The absence of survival in the PBS control group and the high survival among vaccinated animals indicate a strong protective effect under the experimental conditions used.

The rationale for comparing these preparations is based on their complementary antigenic characteristics. Whole-cell inactivated vaccines expose the immune system to a broad array of surface and internal bacterial antigens and retain pathogen-associated molecular patterns that can contribute to innate immune activation. Sonicated preparations, in contrast, provide a complex mixture enriched in soluble and membrane-associated proteins. Because *P. aeruginosa* exhibits considerable antigenic and genetic diversity, multicomponent approaches may have advantages over vaccines directed against a single antigen.

The observed increase in IgG following the first booster indicates a strong secondary humoral response. IgG levels after the second and third immunizations were similar in both vaccine groups, suggesting that the additional booster did not produce a substantial further increase in the measured total IgG under the conditions of this assay. Importantly, similar IgG responses were observed for the two vaccine preparations, supporting comparable immunogenicity based on this endpoint.

Protection following lethal challenge was also broadly comparable. Complete survival was observed in both experiments among mice receiving the formaldehyde-inactivated preparation, whereas survival with the sonicated preparation was 91.7%. The differences between vaccine groups were not statistically significant. Thus, the available data do not support a claim that either preparation is superior in protective efficacy.

SBA provided additional evidence that vaccination induced functional antibodies. The sonicated preparation showed detectable activity at a higher serum dilution than the whole-cell preparation (1:64 versus 1:32). This observation may indicate differences in antibody specificity, avidity, or epitope accessibility; however, the study did not measure these parameters directly. The SBA findings should therefore be interpreted as descriptive evidence of functional activity rather than proof of superior antibody quality.

The findings are consistent with previous experimental studies reporting protective responses following immunization with inactivated *P. aeruginosa* preparations (10,12,13). Earlier work has also demonstrated the feasibility of whole-cell vaccine approaches and highlighted the potential of conserved surface-associated proteins as vaccine targets (14,15).

Several limitations should be considered. First, the experiments were performed in mice and used a relatively small number of animals per group, limiting generalizability to humans and the ability to detect modest differences between formulations. Second, immunogenicity was assessed mainly through total IgG and SBA; cellular immunity, IgG subclasses, mucosal responses, cytokine profiles, memory responses, and other potential correlates of protection were not evaluated. Third, challenge was performed with a single experimental strain, so cross-protection against genetically diverse clinical isolates remains unknown. Fourth, the duration of protective immunity was not determined.

## Conclusions

Formaldehyde-inactivated whole-cell and sonicated protein preparations of *P. aeruginosa* elicited strong antigen-specific IgG responses, generated complement-dependent bactericidal activity, and conferred substantial protection against lethal challenge in mice.

Under the conditions of this study, the two vaccine preparations showed broadly comparable immunogenicity and protective efficacy. These findings support further preclinical investigation of multicomponent *P. aeruginosa* vaccines, particularly studies incorporating multiple clinical strains, cellular and mucosal immune endpoints, optimized formulations, and longer follow-up.

## Acknowledgements

The authors thankfully acknowledge the Department of Microbiology of Dhaka Medical College, Dhaka for providing laboratory support to perform this study and the Departments of Dhaka Medical College and Hospital which permitted sample collection for this study.

This study was funded by Bangladesh Medical Research Council (BMRC) (Ref: BMRC/Research Grant Revenue/2022-2023/34(1–19). However, the funding agency had no role in study design, data collection and analysis, interpretation of results, preparation of manuscript. Decision to submit the manuscript has been obtained from BMRC.

AI tools (unpaid versions of Grammerly and ChatGPT from OpenAI) were used only for English language editing and grammar improvement.

## Supporting information

S1 ARRIVE 2.0 Checklist S2 Human data

